# Stable states in an unstable landscape: microbial resistance at the front line of climate change

**DOI:** 10.1101/2025.02.07.636677

**Authors:** Dylan R. Cronin, Hannah Holland-Moritz, Derek A. Smith, Samuel T. N. Aroney, Suzanne B. Hodgkins, Mikayla Borton, Yueh-Fen Li, Ahmed Zayed, Kieran Healy, Andreas Persson, IsoGenie Field & Analytic Teams 2010-2017, EMERGE Institute Coordinators, Malak M. Tfaily, Patrick Crill, Carmody K. McCalley, Kelly Wrighton, Ruth K. Varner, Gene W. Tyson, Ben J. Woodcroft, Sarah C. Bagby, Jessica Ernakovich, Virginia I. Rich

## Abstract

Microbiome responses to warming may amplify or ameliorate terrestrial carbon loss and thus are a critical unknown in predicting climate outcomes. Because the rapid thaw of permafrost peatlands makes a very large store of soil carbon available to microbial metabolism, understanding microbiome dynamics in these systems is particularly urgent. We quantified microbial warming response over seven years across three habitats in a thawing permafrost peatland, using large-scale multi-omics data. We integrated analyses of organisms (via taxonomy), functions (via metabolic pathways and proteins), and community organization (via network structure and ecological assembly) to deeply characterize response mechanisms. We consistently found a pattern of within-habitat microbiome stability, with virtually no signal of gradual change in the warming period studied. The resistance to change appeared bolstered by habitat-specific dispersal processes and community-level functional redundancy, particularly via versatile carbon generalists. Our findings also reveal key genome-inferred metabolic processes that underlie microbiome stability. Together, our results highlight the importance of understanding the limits of these stabilizing processes and suggest that future research should reorient towards critical habitat transitions.

## Introduction

Arctic permafrost peatlands, warming at a rate four times faster than the global average^1^, contain globally significant reserves of soil organic carbon^2^. This carbon’s fate and flux dynamics as carbon dioxide (CO_2_) or methane (CH_4_) will determine these systems’ feedbacks to climate change^3–5^. Soil microbiota play a critical role in controlling these dynamics (reviewed in Jansson & Hofmockel, 2020), but the community structure, assembly, and metabolism of the microbiota are altered in turn by the cascading changes set off by rising Arctic temperatures^7–9^. Notably, temperature change is expected to directly affect the composition^10^ and interactions^11^ of microbial communities through differential impacts on each microorganism’s metabolic thermodynamics. Alternatively, ecological factors such as functional stability and metabolic flexibility^12,13^ may reinforce community resistance and resilience to gradual change. In either scenario, sufficient disturbance can push communities into non-equilibrium dynamics and trigger reassembly into a new community state^14–16^. Standard operating ranges and baseline stability of microbial communities in a given habitat remain largely unknown^16^ (see Supplementary Note 1). Other major unknowns include uncertainty in the temporal component of a disturbance, timeframe required for response by the microbial community^16^, how variability in soil microbiome structure alters biogeochemical processes^17^ and whether altered ecosystem functioning results from a decoupling between microbial composition and functions^13^. The pressing need to understand how soil microbiomes respond to temperature-induced change has motivated numerous studies^10,18–25^, but to date no clear pattern has emerged. Complicating interpretation, this body of work includes varying study designs and disturbance intensities (observation vs. manipulation^10^), timeframes (few-vs. many-year^21,24^), and environmental settings, in which weather, soil properties, depth, plant communities, and hydrology may outweigh climate warming impact^18–20,22,23,25^(see Supplementary note 2). Despite these challenges, understanding the dynamics of soil microbiome response to warming is vital for establishing how climate models should represent these critical ecosystems^26,27^.

Model ecosystems create an opportunity for integrative study of these dynamics. Stordalen Mire is a discontinuous permafrost peatland in northern Sweden (68° 21’ N, 19° 20’ E; Supplementary note 3) that has been intensively studied over decades as its permafrost thaws. These studies have tracked the collapse of patches of dry elevated palsas (Supplementary Fig. 1) to partially thawed bogs and fully thawed fens, with total fen extent doubling over the past 50 years^5^. Progression along this thaw gradient is marked by both abiotic and biotic shifts affecting redox conditions (predominantly oxic in palsa, anoxic in fen), pH, hydrologic connectivity (dry palsa, perched water tables in bog, permanently inundated fens), and plant, microbial, and viral community composition^5,7,9,28–30^. Importantly, these shifts across the thaw gradient induce increasingly anaerobic microbial lineages and metabolisms, including methanogens^9,29^, driving substantial increases in net radiative forcing^5,31^. Because habitat conversion proceeds in local patches, long-term observation of plant communities has provided many opportunities to observe the conversion of the diverse forbs of the palsa to *Sphagnum*-dominated bog and *Eriophorum*-dominated fen^5,31,32^. These repeated observations suggest that aboveground, on the ecosystem scale, these habitat types behave as alternative stable states^33^, local minima or basins (here meaning more stable) in a stability landscape traversed by communities. Previous work has provided a detailed understanding of microbial communities and processes in short periods within each habitat^9,32,34,35^, but the belowground temporal dynamics within each habitat have so far gone unexplored.

Each of the alternative stable states observed aboveground, in the plant communities at the surface of soil cores, could conceal any of three distinct belowground patterns. These patterns can be visualized in terms of the classic “ball and cup” analog^33^ (Extended Data Fig. 1). First, a habitat’s microbiome could be stable as long as its plant community is stable; a time series of cores with plant communities belonging to one aboveground state would capture only minor fluctuations belowground (Extended Data Fig. 1(i)). Stability could arise either because the community is insensitive to disturbance (resistance) or because it recovers rapidly from disturbance (resilience^12,13,18^). Second, if the microbiome states’ minima are shallow and aboveground-belowground coupling is loose, a microbiome could shift gradually towards the next state in the canonical thaw progression (i.e., palsa to bog, bog to fen) or to a different transitional state (Extended Data Fig. 1(ii.a, ii.b)). Third, a microbiome could shift abruptly between the belowground states associated with palsa, bog, and fen, either leading or lagging aboveground shifts (Extended Data Fig. 1(iii.a, iii.b)). A time series of cores with a given plant community might capture this saltatory microbiome change; however, very long lags could make the plant and microbial communities appear completely uncoupled, while with very short lags the pattern would appear as within-habitat stability, with aboveground state consistently predicting the belowground community. Critically, each of these patterns carries different implications for how predictive modeling should represent a habitat’s soil microbial community and its activity, and for the type and resolution of data required to parameterize models. More broadly, distinguishing between these patterns would reveal the structure of plant-soil feedbacks in a rapidly changing ecosystem.

Here we examined microbiome temporal dynamics in Stordalen Mire’s habitats through three genome-resolved lenses: *organisms* (across taxon ranks ranging from phylum to strain), *functions* (encoded metabolic pathways and proteins), and community *organization* (community network structure and mechanisms of ecological assembly). Our depth-resolved dataset of 199 metagenomes (2.37 Tbp) and 66 metatranscriptomes (1.87 Tbp), from soil cores defined as palsa, bog, or fen by the plant community observed at the surface, captures seven years of progressive ∼0.7°C/yr belowground warming for each habitat (2011-2017, Fig. 1A) at peak growing season. This period included significant interannual fluctuations in air temperature and water table depth (Extended Data Fig. 2) and encompassed four of the planet’s hottest years on record^36^. Through each lens, at all levels of granularity, in each habitat, at nearly every depth, we consistently observed microbiome stability (Supplementary note 1), closely coupled to the alternative stable states aboveground (Extended Data Fig. 1(i)). Together, our analyses suggest that this microbiome stability is driven by (1) metabolic versatility within individual lineages, (2) functional redundancy between lineages, and (3) stochastic dispersal in the waterlogged soils of bog and fen. Moreover, our integrative approach suggests new hypotheses on nutrient-cycling pathways and partnerships in each of Stordalen Mire’s habitats.

**Figure 1:**
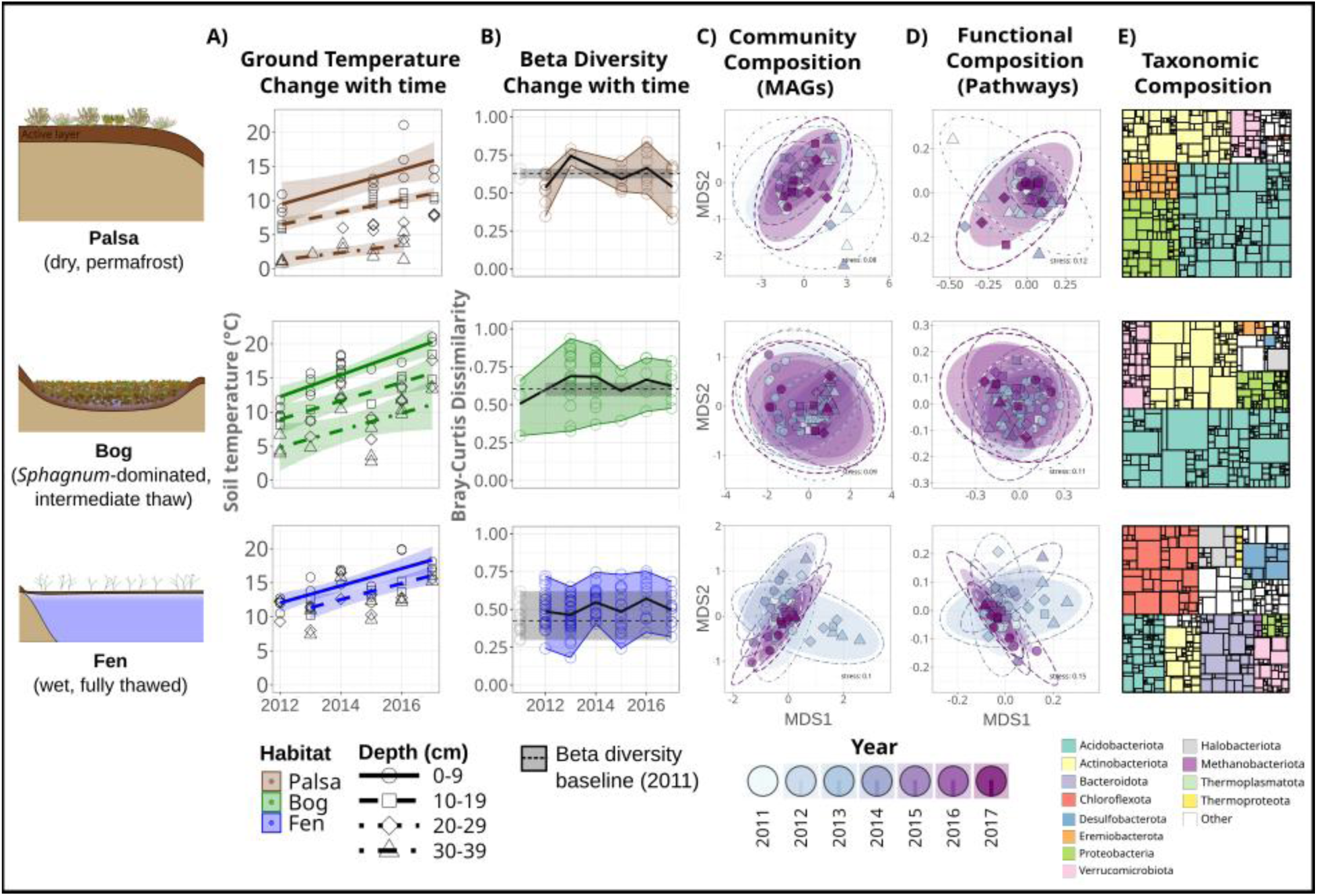
Stordalen Mire microbiomes resist temporal change. **A)** The three main thaw-gradient habitats (palsas, bogs and fens; colored throughout as brown, green and blue, respectively) exhibit significantly increasing soil temperatures with time (lines indicate significant linear relationships within each depth, p<0.05). Shapes of points correspond to depths; 0-9: circle, 10-19: square, 20-29: diamond, 30-39: triangle. **B)** Microbiome beta diversity did not change significantly with time. Points are pairwise comparisons of Bray-Curtis dissimilarity to samples at the same depth against the earliest baseline year for which at least two depth-matched microbiomes were available, and black lines are the running dissimilarity average. Only 0-9 cm are shown. Reference-year microbiomes’ within-year dissimilarities are indicated by gray dots, with their ranges and averages extended as shaded gray bars and dashed lines, respectively. **C,D)** Habitat and depth structure microbiome beta diversity data more strongly than year, observed in non-metric multidimensional scaling (NMDS) ordinations of each sample’s MAG-based taxonomic **(C)** and functional **(D)** composition colored by year (purple shading). Depths are denoted by shape (as in **(A)**) and demarcated by ellipses (dashed-line ellipses reflect each habitat’s multivariate normal distribution, while shaded ellipses are the multivariate t-distribution). Functions are cumulative pathway abundances: each complete pathway observed in a MAG was weighted by that MAG’s relative abundance, then all occurrences are summed for each pathway. **E)** Overall taxonomic composition per habitat (treemap with dereplicated MAGs as rectangles, colored by phyla and sized proportional to average relative abundance over time).

### Microbiomes are resistant to increasing ground temperature

#### Microbiome composition and function resisted change over time

To evaluate microbiome changes, we reconstructed 13,290 metagenome-assembled genomes (MAGs) that represent 1,806 species-level taxa (i.e., clustered at 95% average nucleotide identity, Fig. 1E). Together these MAGs captured 73% of the sequenced community at genus-level [see Methods], which is high for natural systems. Read-mapping-based abundances revealed that microbial diversity and composition were resistant to change over seven years within habitat and depth (down to 39 cm) (Fig. 1A-E, Supplementary Fig. 2, Supplementary Table 1). Very little variation in community composition could be accounted for by sample year at any level of resolution (strain: ≤3.0%, p = 0.03); genus: 1.6%, p = 0.03; phylum: 1.6%, p = 0.005) evaluated by distance-based redundancy analyses on single-copy marker genes (Fig. 2). Similarly, in 92% of habitats and depths, genome-resolved community composition was not explained by time (Fig. 1B-D, Fig. 2, Extended Data Fig. 3) and inter-annual fluctuation was significant only at certain depths in the fen (for 0-9, 20-29, and 30-39 cm, respectively: R^2^: 0.35, 0.62, 0.81; p-value: 0.036, 0.048, 0.048; Extended Data Fig. 4A). Only the deeper fen (30-39 cm) samples exhibited both significant inter-annual fluctuation (Extended Data Fig. 3, Extended Data Fig. 4A) and directional change (R^2^ = 0.50, p = 0.012) in community composition with time (Fig. 2), and only 8 of 1,806 species were significantly correlated to time (Spearman rho ranging from -0.7 to 0.91, p < 0.05; Extended Data Fig. 4B-C) (see Supplementary note 4). The stability observed across a range of taxonomic granularities suggests that the microbial community is compositionally resistant over time.

**Figure 2:**
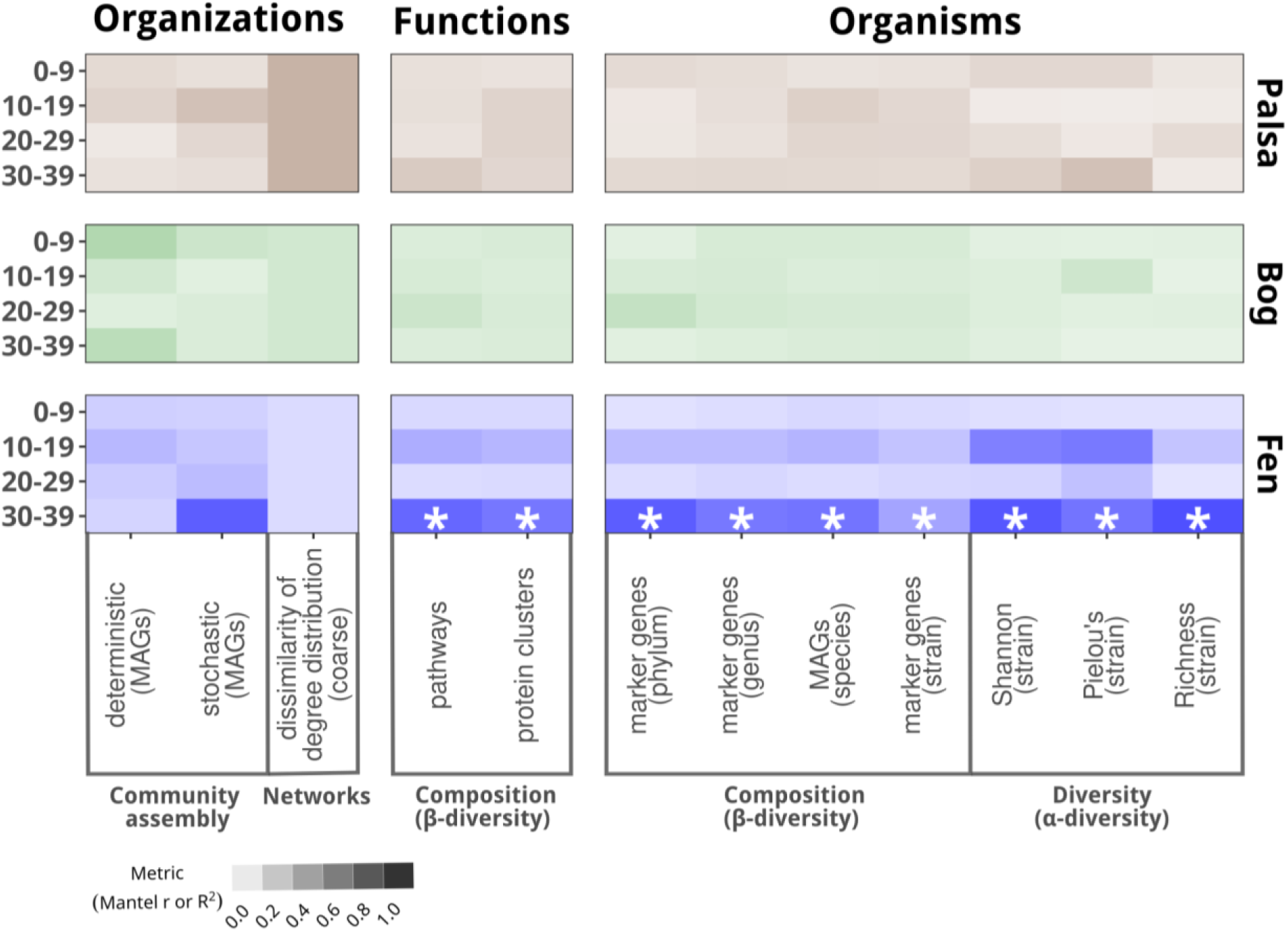
Heatmap of microbiome temporal change in each habitat and depth demonstrating that organizations, functions, and organisms exhibit temporal stability across nearly all Stordalen Mire habitats and depths examined. Change was assessed spanning **organizations** (both community assembly metrics of stochasticity and determinism and network structure), **functions** (from pathways to protein clusters, both annotatable and non-annotatable), and **organisms** (from phylum to strains diversity metrics, both single copy marker genes and MAGs). The amount of variation explained by time (as Mantel R or R^2^ depending on statistical test appropriate for each data type; see Methods) is indicated with the intensity of the shaded color; white asterisks denote significant (p<0.05) correlations to time. Across all metrics surveyed, only deeper fen (30-39 cm) showed significant change over time.

Encoded microbiome functions were similarly consistent within habitats over time (Fig. 1D, Fig. 2, Fig. 3A). Beta diversity of community-wide protein clusters (a metric unconstrained by functional annotation), as well as manually-curated genome-resolved pathways of carbon, nitrogen, and sulfur metabolism did not significantly change over time within 92% of habitats and depths (Fig. 2, Fig. 3, Extended Data Fig. 5); the sole exception was 30-39 cm in the fen (R^2^: 0.49-0.55, PERMANOVA, p < 0.05 Fig. 1C-D; Fig. 2; Extended Data Fig. 4A). At the coarsest level of functional annotation (MAG-specific specialization), individual functional guilds of organisms (e.g., methanogens, homoacetogens, fermenters, and macromolecule degraders) maintained similar abundances through time. (Expressed microbiome function was not used for temporal analysis due to discontinuity in the metatranscriptomic time series.) Thus, within the majority of Stordalen Mire habitats and depths, we observed both compositional and functional resistance to change over time across multiple levels of granularity.

**Figure 3:**
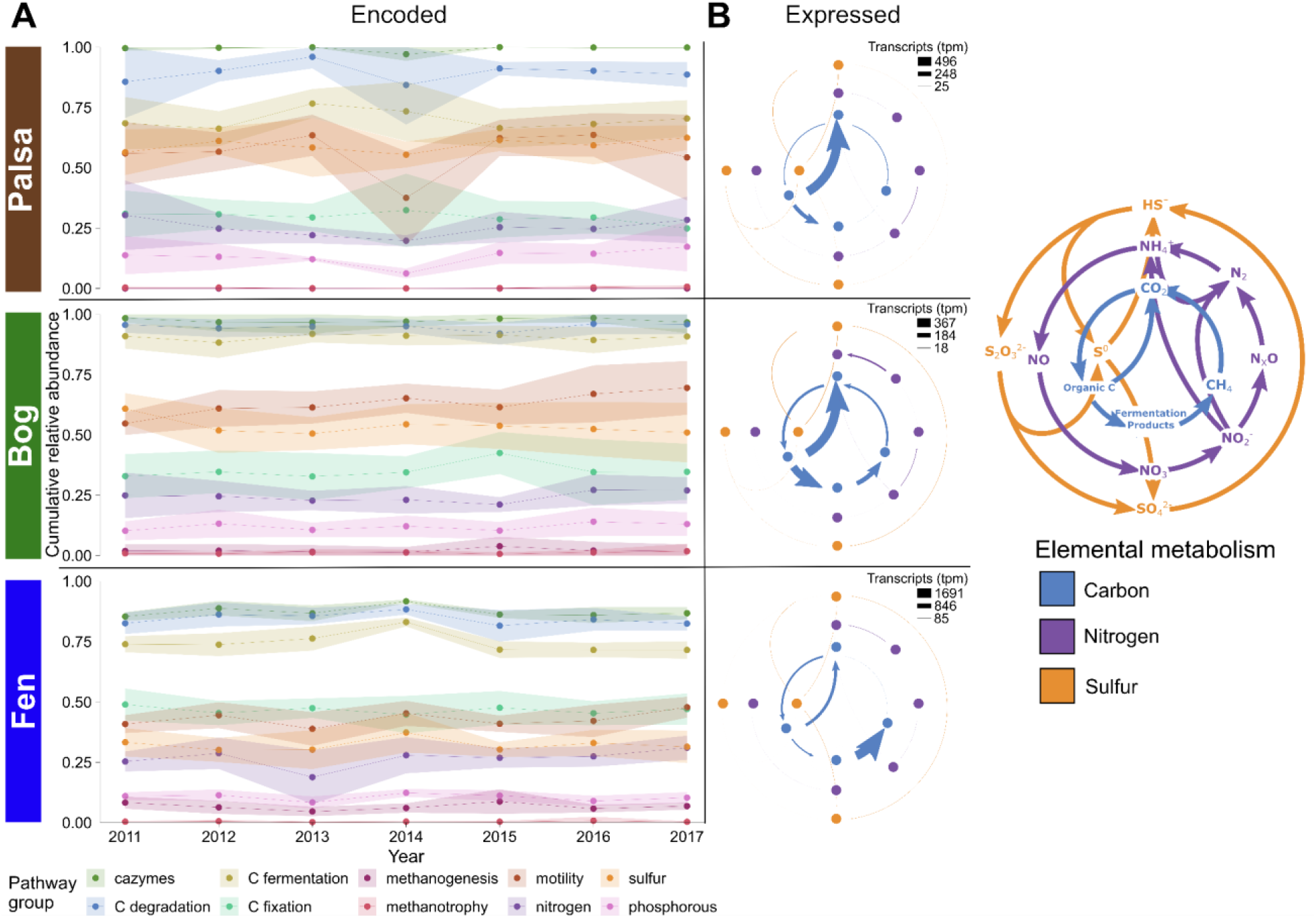
Microbiome metabolisms are stable over time but differ across the thaw gradient. **A)** MAG-encoded cumulative relative abundance of carbon, nitrogen, sulfur and phosphorous metabolic pathways over time. Points and ribbons represent mean ± standard deviation across depths and replicates. **B)** Aggregated over the study period, key pathways in carbon, nitrogen, and sulfur metabolism from metatranscriptomic data show differences in microbial investment in methanotrophy and carbon fixation, nitrogen fixation, and sulfur redox cycling. A reference figure showing the transformations mapped is shown on the right.

Our third lens, microbial community organization, also revealed stability over time within each habitat and depth. We used two distinct genome-resolved network approaches to understand the influence of soil biogeochemical conditions (Weighted Genome Correlation Network Analysis (WGCNA)) and inter-microbe interactions (multilayer temporal networks (MTN)) on the organizational structure of the community with time. Habitat-specific WGCNA modules showed no linear correlation of community composition to time (Pearson, p<0.05) (Supplementary Table 2). MTN microbial co-occurrence networks exhibited variability in specific microbial interactions over time (Fig. 4A, Extended Data Fig. 6A-C; Jaccard similarity coefficient of triangles), though the overall network structure remained more consistent at broader scales (Fig. 2, Fig. 4B, Extended Data Fig. 6A-C; degree distribution and degree correlation). This suggests that broader community network patterns are resistant to change, even as individual interactions shift temporally. To understand if subcommunities of microbes were resistant over time, we asked how many years each MTN subcommunity persisted (Extended Data Fig. 6D). Persistent subcommunities (occurring in ≥ four of the seven years) accounted for a conservative estimate of 40% of the abundance-weighted community with time (depending on the interannual relax rate; Extended data Fig. 6E); thus, while not all microbe-microbe interactions were stable from year to year, persistent clusters contributed extensively to community composition.

**Figure 4:**
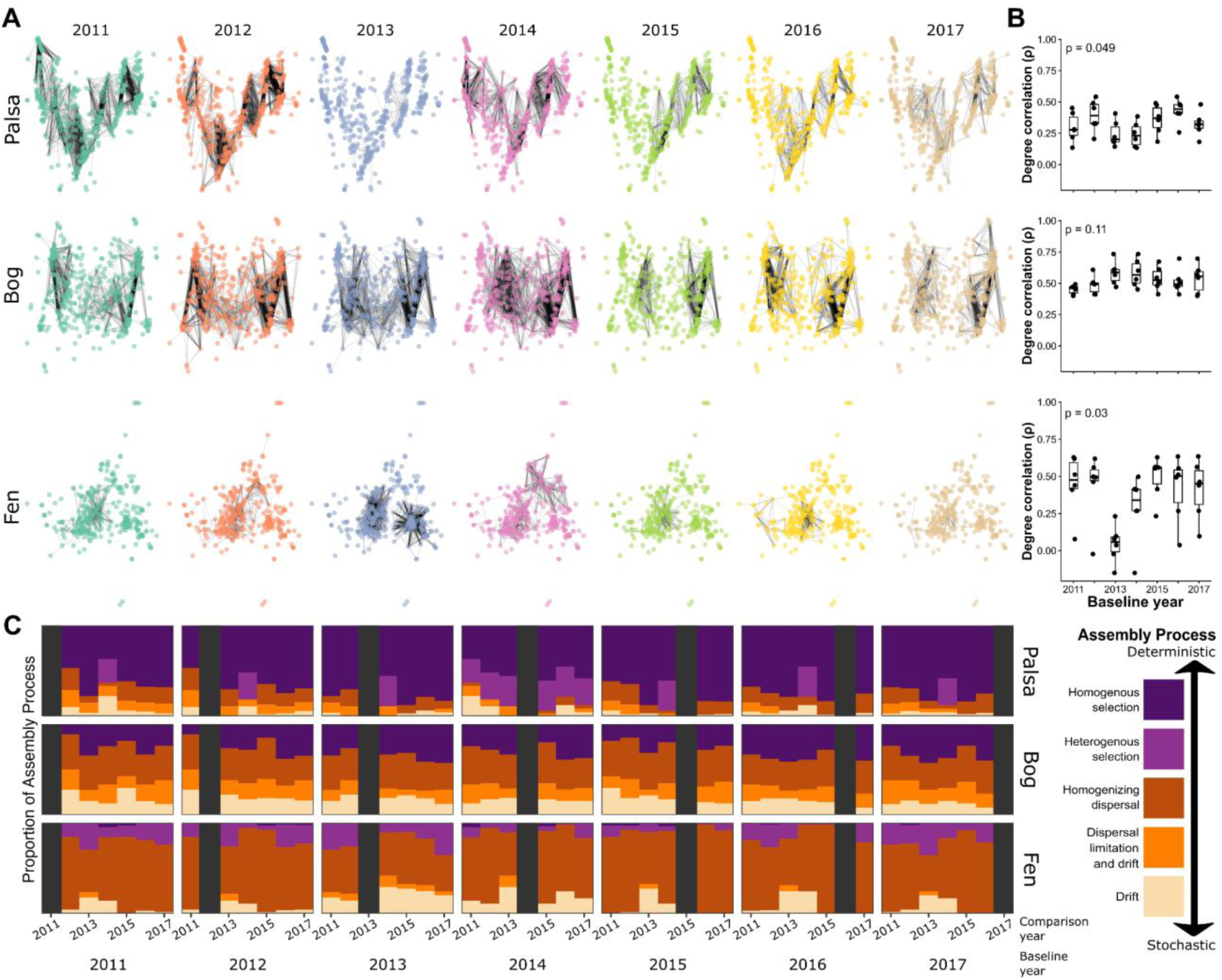
Microbiome organization is consistent within habitats over time. **A)** Multilayer temporal networks (MTN) depict correlations between MAGs within annual networks for each habitat. Each network layer corresponds to a specific year, with nodes representing MAGs. Black edges represent significant (p<0.05) correlations (>0.3) between MAGs within the same habitat and year. **B)** Pairwise interyear correlations (Spearman’s rho) in network node degree (x-axis, base year; n=6 interyear comparisons per baseline year; non-parametric Kruskal-Wallis test) within each habitat. Top, palsa; middle, bog; bottom, fen. Boxplots show the median (center line), the interquartile range (box), and whiskers drawn to 1.5× the IQR. Data points beyond the whiskers are shown as outliers. Note that the fen’s recovery to a near-stable state after 2013 suggests strong resilience in this habitat. **C)** Assembly processes detected within each habitat (rows) by β-NTI and RCBC modeling in interyear comparisons. For each year, columns show pairwise comparisons to all other years (self-comparisons are obscured by black bars). Stacked bar colors denote the proportion of different assembly processes detected in each pairwise comparison, revealing overall stability in assembly within habitats over time.

Finally, we sought to understand whether the ecological mechanisms that reinforce stability in each habitat’s community organization were themselves changing over time. Constructing null models with randomization procedures informed by phylogenetic and taxonomic community composition [see Methods], we used beta-null modeling^37^ to determine whether stochastic (dispersal and ecological drift) or deterministic (selection) processes drove community structure in each habitat (Fig. 4C, Extended Data Fig. 7). A change in community assembly mechanisms over time could indicate a shift in the ecological pressures shaping the community (for example, increased selection due to changing ground temperature). Instead, beta-null modeling showed that while each habitat was governed by distinct assembly mechanisms (homogenizing selection in the palsa, homogenizing dispersal in the water-logged habitats), the ratio between deterministic and stochastic assembly processes within each habitat did not change with time (Mantel, p > 0.05) (Fig. 2, Fig. 4C).

#### Microbiome composition and function had weak susceptibility to temperature

Finally, we explored the possibility that organisms, functions, and organization of the communities were being shaped by subtle or non-monotonic changes in abiotic drivers. Below we focus on temperature, since other abiotic factors like pH and redox-proxies did not change with time (Supplementary Fig. 3). After accounting for covariation between temperature and depth (strongest in palsa; Extended Data Fig. 2), genome-resolved community composition was slightly, but significantly, linearly correlated to ground temperature in bog and fen but not palsa (Fig. 2, Bray-Curtis dissimilarity: partial mantel rho = 0.152 and 0.24, p < 0.001, in bog and fen respectively). However, when habitat and depth were included in non-linear generalized additive models, no predictive relationship remained between community composition and temperature (p = 0.4; Supplementary note 5), suggesting that the effect of temperature was weak. Echoing these results, while some WGCNA modules were significantly (Pearson, p < 0.05) correlated with temperature, all but one of these modules were more strongly correlated with depth. Some of these WGCNA modules were more strongly correlated to carbon cycling metrics (e.g., ẟ^13^C_CO2_, ẟ^13^C_CH4_, ɑ_CO2-CH4_ (Extended Data Fig. 8), consistent with previous work at the site^9,11,29,30,38^, and other features (e.g., pH and total nitrogen). Ground temperature had no significant impact on carbon, nitrogen, or sulfur pathways in the palsa or bog, and only on xyloglucan degradation in the fen (Supplementary note 5). No community assembly mechanisms were related to ground temperature or air temperature (day-of-sampling or prior four weeks) (Mantel, p > 0.05), implying that even with lag effects, temperature was not a strong selective or stochastic force on community formation. As with temporal change, microbiome sensitivity to temperature was strongest in the fen but weak overall and overshadowed by other abiotic factors (depth, pH, and nutrient availability).

### Metabolic connections maintain stability

We next asked whether and how the activity of the microbiota within these communities might contribute to reinforcing within-habitat stability. Microbiome capacity to degrade organic matter depends not only on organic matter availability but also on access to other nutrients including bioavailable nitrogen and sulfur. Access to nitrogen is a fundamental constraint on microbial growth in northern peatlands, where nitrogen is frequently limiting^39,40^. Similarly, despite small environmental pools of sulfur species^38,41^, sulfur cycling can control organic matter degradation rates, particularly in the bog^34^. Bottlenecks in nitrogen, sulfur, or carbon flow due to warming-dependent changes in organic matter profiles, redox conditions, or enzyme kinetic^421^ might disturb the community, but access to alternative pathways or partners for substrate processing could confer resistance to such pressure.

#### Chitin degradation and nitrogen fixation offer alternative pathways to nitrogen acquisition

When preferred low-molecular-weight organic nitrogen compounds^43,44^ are not available, nitrogen fixation and chitin degradation can provide alternative nitrogen sources. Nitrogen fixation increases the bioavailable nitrogen pool, benefiting diazotrophs directly and the community indirectly^45^. Up to 25% of the Stordalen Mire community encoded nitrogen fixation (compared to ∼0.3% in marine microbiome^46^), and > 75% of encoding MAGs showed pathway expression (Extended Data Fig. 9E).

Pathways for degrading chitin ((C_8_H_13_O_5_N)_n_), the second most abundant biopolymer in nature^47^, were even more widespread. An insoluble compound whose breakdown requires extracellular enzymes, chitin is targeted by many degradation pathways. At Stordalen Mire, multiple chitin degradation pathways were ubiquitously encoded (abundance weighted median 73%, range 45%–94% (Extended Data Fig. 9F)) and expressed (median 95% of encoding cells, range 59–100% with one outlier sample at 5%). Furthermore, organisms containing these functions were closely linked to key environmental outputs in WGCNA networks (as Variable Importance in Projection (VIP) members of modules related to carbon and nitrogen cycling (Extended Data Fig. 8)). In many aquatic and terrestrial environments, including peatlands, extracellular chitin degradation products can facilitate commensal and cross-feeding interactions^47–50^, which may promote community stability^51^ while buffering against fluctuations in low-molecular-weight nitrogen availability. The ubiquity of both nitrogen fixation and chitin degradation pathways across Stordalen Mire habitats may likewise confer resistance.

#### Cryptic sulfur cycling and degradation of sulfur-containing polysaccharides might minimize sulfur limitations

Cryptic sulfur cycling can profoundly influence system biogeochemistry^52^. We observed habitat-wide expression of dissimilatory sulfur oxidation and sulfate reduction (encoded pathway relative abundances ∼25% and up to 10%, respectively (Extended Data Fig. 9I,J), suggesting the active use of cryptic sulfur cycling at Stordalen Mire. Interestingly, although all microbes require sulfur for growth, we detected assimilatory sulfate reduction in only 25% (fen) to 50% (palsa, bog) of the population (Extended Data Fig. 9K). Our analysis of carbohydrate degradation suggests a novel hypothesis: community members may meet some of their sulfur requirements through CAZyme degradation of sulfur-containing polysaccharides (Extended Data Fig. 9L), known to be preferentially consumed from plant organic matter^38^. This novel linkage between the carbon and sulfur cycles in permafrost peatlands is analogous to some marine isolates’ ability to degrade sulfur polysaccharides and thereby increase H_2_, acetate, and sulfate concentrations^53,54^. Such a pathway is expected to release sulfur extracellularly, which could serve as a public good and partially alleviate sulfur limitation for other members of the community.

#### Carbon generalists are highly prevalent, and macromolecule degradation is widespread

We defined carbon generalists as MAGs encoding ≥ 3 alternative CAZyme pathways and ≥ 3 sugar degradation pathways (Supplementary note 6; Extended Data Fig. 10). These sorts of acquisition-oriented carbon generalists are typical of cold peatlands and other wetlands^13,55,56^. In Stordalen Mire, such MAGs constituted >80% of the community and were taxonomically diverse in each habitat (Fig. 5A) and at nearly all depths. The prevalence of these metabolic generalists produced community-level functional redundancy, with each CAZyme pathway encoded by a median of 531 carbon generalists, and each sugar degradation pathway by a median of 253. Macromolecule and sugar degradation dominated encoded and expressed pathways, particularly in the palsa and bog where >84% of the abundance-weighted community encoded and >50% expressed these pathways (Fig. 3A; Extended Data Fig. 5). Fermentation pathways were also highly abundant (47-93% cumulative relative abundance; Fig. 3A; Extended Data Fig. 5). Collectively, functionally redundant carbon generalists should stabilize the flux of carbon through the system to downstream partners even as environmental conditions and carbon inputs shift.

**Figure 5:**
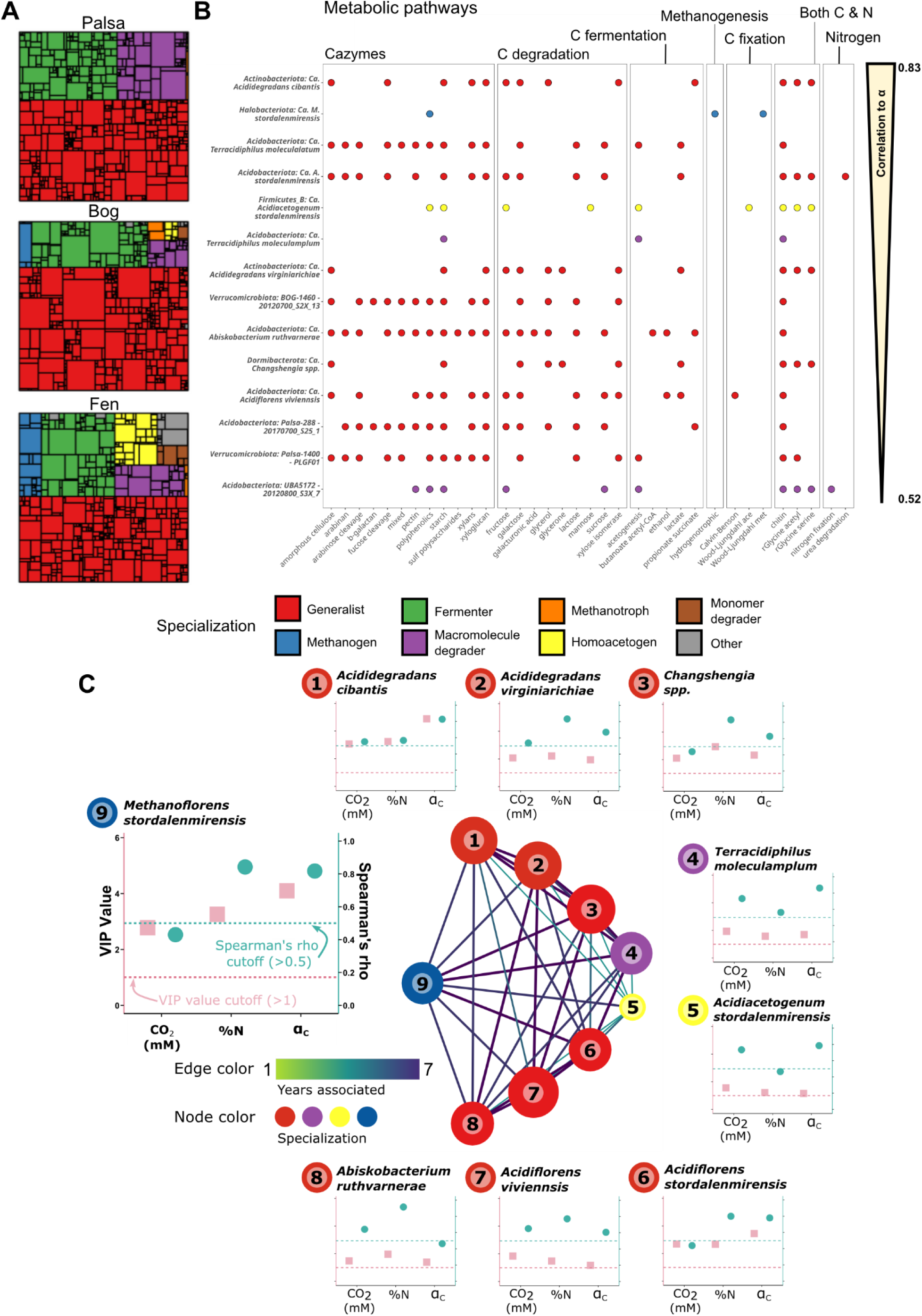
Microbiome metabolisms across the thaw gradient are interlinked by carbon and nitrogen cycling. **A)** Functional composition of palsa, bog, and fen communities. Treemap rectangles are MAGs colored by carbon metabolic specialization and scaled by relative abundance. **B)** Carbon and nitrogen-related pathways present in MAGs belonging to the weighted genome correlation network analysis (WGCNA) module correlated with the isotopic fractionation of carbon between carbon dioxide and methane carbon (ɑ_CO2-CH4_) in the bog. MAGs are colored by carbon metabolic specialization as in Fig. 3 and ordered by VIP score. **C)** Bog network structure of the persistent subcommunity of MAGs connected to *Ca*. ‘M. stordalenmirensis’ in multi-layer temporal network (MTN) modeling (n=7). MAGs were included if they maintained a connection to *Ca.* ‘M. stordalenmirensis’ for at least six years, had a VIP score > 1, and a Spearman correlation > 0.5 to at least one environmental variable. Edge color represents the number of years adjoining MAGs were connected; node color indicates MAG specialization, with colors as in (A). We note that many other MAGs were connected to the methanogen, but not persistently. Subplots show the Spearman correlation coefficient (rho) of each MAGs to key environmental variables (teal) and their VIP value (pink).

#### The flexible community of carbon generalists supports methanogenesis in the bog and fen

Despite low abundances, methanogens have an outsized impact on ecosystem output, creating a pressing need to understand the mechanisms that support their activity. Consistent with previous reports^9^, methanogenesis—primarily hydrogenotrophic—was encoded by a handful of MAGs (< 1%) in the bog and fen and was highly expressed there (Fig. 3). Acetoclastic and methylotrophic methanogenesis were uncommon, with the former present only in the fen (Extended Data Fig. 5). We observed that several methanogens persisted throughout our study period (Fig. 5B-C, Supplementary note 7), even though the activity of these metabolic specialists depends on the supply of a narrow range of substrates. Diverse metabolic partnerships that reduce substrate limitation for methanogens have been observed to stabilize aquatic microbial communities^57,58^. We asked whether bog and fen methanogens might likewise partner with generalists and leverage these partners’ metabolic flexibility to buffer thaw-induced change.

Our network analyses allowed us to address this question in two ways. First, we examined WGCNA relationships to the fractionation factor ɑ_CO2-CH4_, an isotopic indicator of CH_4_ metabolic pathways. The hydrogenotrophic methanogen *Ca.* ‘Methanoflorens’ spp. has previously been observed to be a strong predictor of ɑ_CO2-CH4_^9,29^. Strikingly, WGCNA not only supported the correlation of *Ca.* ‘Methanoflorens’ spp. with ɑ_CO2-CH4_ but also identified thirteen non-methanogens (including multiple *Acidimicrobiales*, *Terracidiphilus* and *Ca*. Acidiflorens) that were highly predictive of (VIP > 1.5) and correlated to ɑ_CO2-CH4_ (Spearman’s rho, p < 0.005). These correlations were comparable to or higher than that of *Ca.* ‘M. stordalenmirensis’ itself (Fig. 5B), suggesting that these predominantly carbon-generalist, nitrogen-flexible lineages may be gatekeepers for methanogenic carbon flux. For example, several of the *Ca*. ‘Acidiflorens’ spp. may supply H_2_ and formate to *Ca.* ‘M. stordalenmirensis’^9^. Second, we identified persistent MTN associations between *Ca*. ‘M. stordalenmirensis’ and with eight MAGs—all VIPs for ɑ_CO2-CH4_, [CO_2_], ẟ^13^C_CO2_, and/or nitrogen content—from the same WGCNA module (Fig. 5C). These MAGs included the Dormibacterota *Ca*. ‘Changshengia’ spp., which was previously identified as highly correlated with the fraction of carbon mineralized to CO_2_ vs. CH_4_ in the bog^9^.

From close analysis of the metabolic capabilities of methanogen-associated lineages, we propose a novel hypothesis that trading and acquisition of both nitrogen and carbon underlies these relationships. We observed that, whereas methanogens with few network connections tend to encode and express nitrogen fixation, highly connected methanogens important to network structure (e.g., *Ca*. ‘M. stordalenmirensis’) typically lacked recognizable nitrogen-fixation or other nitrogen-linked decomposition pathways (Supplementary note 7, Supplementary Fig. 4, Supplementary Fig. 5). In other systems, methanogenesis expression^59^ and CO_2_ and CH_4_ production increase with addition of chitin, which methanogens do not themselves degrade^49,60,61^. Here, all eight persistent methanogen partners (Fig. 5C) encode and express chitin degradation, potentially mitigating nitrogen limitation for methanogens. Furthermore, we identify a potential new linkage between methanogens and their partners through the reductive glycine pathway. This energy-conserving and amine-recycling carbon-fixation pathway, capable of producing methanogenesis substrates (pyruvate, acetate, and formate) and bioavailable nitrogen (as ammonia^62,63^), is encoded and expressed by five of the eight methanogen partners. These co-occurrence-based observations suggest that nitrogen may play a previously unrecognized role in metabolic interactions supporting methanogen persistence and activity in these ecosystems.

### Habitat-specific microbiome resistance arises from functional redundancy and dispersal mediated by hydrologic connectivity

Stordalen Mire is warming, causing rapid permafrost thaw and habitat conversion^5^ between visibly different alternative stable states aboveground. Our analysis of the soil microbiome’s organization, organisms, and functions in cores defined by their surface plant communities suggests that belowground communities in palsa, bog, and fen are likewise stable and tightly coupled to the aboveground state, to within the limits of our study’s ability to detect lags. What ecological dynamics maintain the microbial communities’ resistance to change? We hypothesize that functional redundancy, metabolic versatility, and homogenizing dispersal drive ecological resistance (Fig. 6).

**Figure 6:**
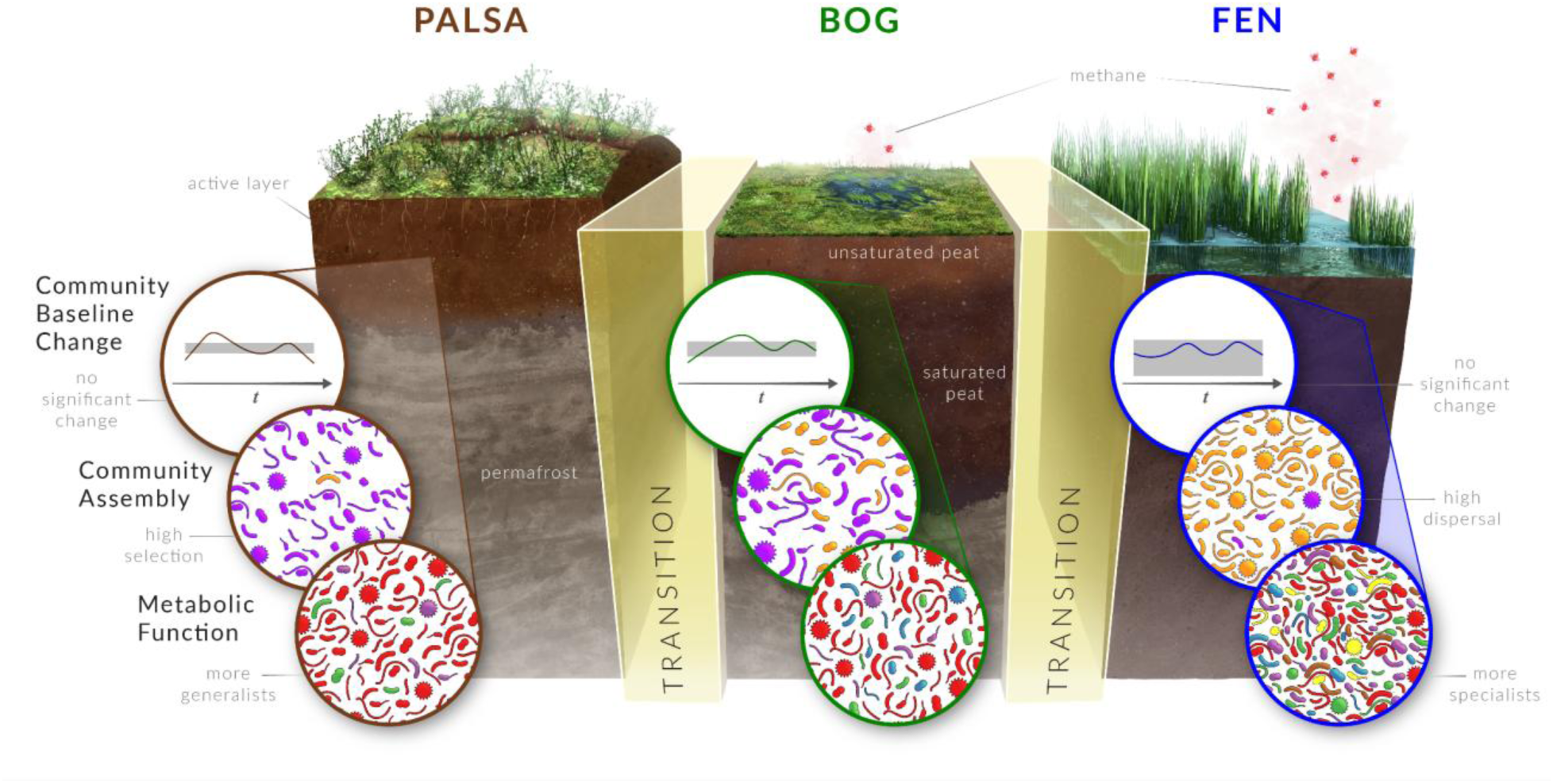
Within-habitat, communities resist change until increasing environmental pressures trigger transitions along the thaw gradient. Community composition fluctuates around a mean but does not change steadily with time in each habitat. Population density increases in tandem with soil saturation^9^. Across the thaw gradient, ecological assembly shifts from predominantly deterministic processes in the palsa (community assembly, purple) to predominantly stochastic processes in the fen (orange), while the balance of functional specializations shifts from predominantly generalists (red) to a more metabolically diverse range of specialists. We hypothesize that functional redundancy, dispersal (bog, fen), and metabolic versatility (palsa, bog) drive within-habitat ecological resistance. Dividers marked “transition” represent the as-yet unknown set of processes and transformations that lead to habitat transition.

Functional redundancy and functional similarity have been proposed to stabilize microbial communities via the “portfolio effect”^64–66^. We saw strong signals of both community-level functional redundancy and species-level metabolic versatility in the stable microbiomes of palsa, bog, and fen, each of which was dominated by diverse taxa encoding multiple alternative pathways for carbon and nitrogen acquisition (Fig. 3, Fig. 5, Supplementary note 6). Studies of generalists have typically focused on habitat generalists rather than resource generalists^67^ (Supplementary note 6), leaving the prevalence of versatile resource generalists in most habitats unknown. We argue that resource generalists merit attention as ecological stabilizers due to their potential for a species-level portfolio effect. That is, just as community function can be maintained by reciprocal changes in efficiency between two compensating species, so should a metabolically versatile cell be able to maintain metabolic activity by rebalancing flux between alternative pathways as conditions change. Increased access via mobile genetic elements to genes providing metabolic versatility would further backstop these stabilizing forces^68^. If redundant pathways within redundant populations stabilize communities, more narrowly distributed specialists^13^ would be shielded from disturbance and the loss of taxa from critical functional groups.

Bog and fen microbiomes are likely further stabilized by homogenizing dispersal, i.e., dispersal high enough to result in a community that is more similar to a neighbor than would be expected by chance (Fig. 4C, Fig. 6, Extended Data Fig. 7). Dispersal dynamics are particularly important during disturbance, when dispersing organisms can determine ecosystem trajectory^69^. Local dispersal can promote community resistance and resilience by “rescuing” disturbed communities, preventing cross-site divergence after local population reductions or extinctions^12,16^. This is likely synergistic with functional redundancy and metabolic versatility, which improve the odds that dispersal will resupply organisms that fill key roles.

### Conclusions

At Stordalen Mire, aboveground plant community reassembly makes habitat conversions detectable from space^70^, but it is the belowground community that drives methane flux, and the dynamics of belowground change are unknown and not easy to remotely observe. Such knowledge gaps are a challenge in eco- and earth system climate models, which must calibrate the representation of microbial complexity^26^ often in the absence of high-resolution datasets and parameter estimation^71,72^. The analyses presented here demonstrate that, despite warming (Fig. 1, Extended Data Fig. 2), landscape-scale change^5^ (Supplementary Fig. 1A), and increasing methane emissions (Supplementary Fig. 1B), habitat-specific microbiome complexity can be distilled to essential functions that are stable during peak growing season on a near-decadal timescale.

Collectively, our findings offer new insights into processes underlying microbial community stability in critical permafrost ecosystems. Our integrated analysis over multiple levels of granularity for organismal ranks, metabolic functions, and community organization revealed little to no significant temporal change within habitats over seven years of progressive belowground warming. This analysis establishes a collection of metrics for assessing community response to environmental disturbance. Multiple independent analyses suggest that the stability of palsa, bog, and fen microbiomes under temperature-induced climate change depends on functional redundancy within communities and dispersal-mediated resupply of organisms encoding essential metabolic versatility.

To complement these insights into Stordalen Mire’s microbiome stability landscape, future sampling must target periods of habitat transition (Fig. 6), when stabilizing capacity is exceeded. Further detailed study at spatial, temporal, and genetic resolutions not captured in this study may reveal other mechanisms of microbial response to changing conditions, as well as improving bounds on any lag between aboveground and belowground stable-state changes. For instance, spring and fall observations capturing changing phenology may illuminate an erosion of stability driven by aboveground/belowground asynchronies. In addition, increased application of long-read sequencing may help resolve strain-level functional differences, including those mediated by the activity of mobile genetic elements. Despite these remaining unknowns, this work reveals the clear existence of stable microbial communities from habitats in a rapidly warming unstable Arctic landscape.

## Collective Author Designations

### IsoGenie Field and Analytic Teams 2010-2017

Jeffrey P. Chanton¹, Darya Anderson², Gareth Trubl³, Joachim Jansen⁴, Moira Hough⁵, Nicole Irwin-Raab³, Rachel M. Wilson¹, Rhiannon Mondav⁶, Robert Jones³, Sky Dominguez⁷, Apryl Perry⁸

1. Earth, Ocean, and Atmospheric Sciences, Florida State University, Tallahassee, FL, USA
2. Department of Soil, Water, and Environmental Sciences, The University of Arizona, Tucson, AZ, USA
3. Department of Microbiology, The Ohio State University, Columbus, OH, USA
4. Department of Ecology and Genetics, Uppsala University, Uppsala, SE
5. College of Forest Resources and Environmental Science, Michigan Technological University, Houghton, MI, USA
6. Centre for Environmental and Climate Science, Lund University, Lund, SE
7. Department of Ecology and Evolutionary Biology, The University of Arizona, Tucson, AZ, USA
8. Department of Earth Sciences and Earth Systems Research Center, University of New Hampshire, Durham, NH, USA

### EMERGE Institute Coordinators

Jeffrey P. Chanton¹, Matthew B. Sullivan^2,3^, Regis Ferriere⁴, Scott R. Saleska⁴, Rachel M. Wilson¹, Benjamin Bolduc², Eoin L. Brodie^5^, Maria Florencia Fahnestock^6^, Michael Ibba^7^

1. Earth, Ocean, and Atmospheric Sciences, Florida State University, Tallahassee, FL, USA
2. Department of Microbiology, The Ohio State University, Columbus, OH, USA
3. Center of Microbiome Science, The Ohio State University, Columbus, OH, USA
4. Department of Ecology and Evolutionary Biology, The University of Arizona, Tucson, AZ, USA
5. Earth and Environmental Sciences, Lawrence Berkeley National Laboratory, Berkeley, CA, USA
6. Department of Natural Resources and the Environment, University of New Hampshire, Durham, NH, USA
7. Schmid College of Science and Technology, Chapman University, Orange, CA, USA

## Non-author contributions

This study has been made possible by data provided by Abisko Scientific Research Station and the Swedish Infrastructure for Ecosystem Science (SITES) as well as the Swedish Polar Research Secretariat. We thank the Swedish Polar Research Secretariat and SITES for the support of the work done at the Abisko Scientific Research Station. A portion of this analysis and non-monetary technical support was also provided through the U.S. Department of Energy user facilities programs. Computational work was supported in part by an allocation of computing time from the Ohio Supercomputer Center. Lastly, we thank contracted science artist Elena Hartley for co-development of Fig. 6.

## Author Contributions

DC, SA, DS, and HH-M all contributed equally to this work. Co-first authors have the right to list their name first in their CV. In addition, VR, SB, GT, BW, and JE are all co-senior authors and contributed equally to this work, order was paired with first author order and all may list their name last on their CV. Following the CRediT model, authors made the following contributions

Conceptualization - VR, CM, SH, RV, EMERGE Institute Coordinators, DC, HH-M, DS, SA, JE, SB, BW, MT, PC

Data Curation - DC, HH-M, DS, SA, VR, BW, SH, FL, RV

Formal Analysis - DC, HH-M, DS, SA, CM, KW, MB, IsoGenie Field & Analytic Teams 2011-2017

Funding acquisition - VR, JE, SB, BW, GT, KW, SH, RV, EMERGE Institute Coordinators, MT, PC

Investigation - DC, HH-M, DS, SA, CM, SH, FL, RV, IsoGenie Field & Analytic Teams 2011-2017, PC

Methodology Development - DC, HH-M, DS, SA, VR, PC

Project Administration - DC, HH-M, DS, SA, VR, JE, SB, BW, GT, KW, SH, RV, EMERGE

Institute Coordinators, PC

Resources Provision - VR, KW, SH, RV, EMERGE Institute Coordinators, PC

Software Programming - DC, HH-M, SA, KW, MB, SH

Supervision Oversight - VR, JE, KW, RV, MT

Validation Verification- SA, DC, HH-M

Visualization - DC, HH-M, DS, SA, VR, SB, KH

Writing - DC, HH-M, DS, SA, VR, JE, SB, BW, GT, CM, KW, MB, SH, FL, RV, IsoGenie

Field & Analytic Teams 2011-2017, EMERGE Institute Coordinators, KH, MT, PC

## Data and materials availability

Sample Metadata Sheet v1.0.0, containing field sample information and geochemical measurements are available on Zenodo at https://zenodo.org/records/12827096. Gas flux measurements are available on Zenodo at https://doi.org/10.5281/zenodo.13377109 and https://emerge-db.asc.ohio-state.edu/datasources/0138_DailyFlux_20122018. Temperature and water table summary for lagged effects analysis are available on Zenodo at https://doi.org/10.5281/zenodo.14189390. Gas flux summary for lagged effects analysis is available on Zenodo at https://doi.org/10.5281/zenodo.14834760. Water table depth measurements used for determining lagged effects are available on Zenodo at https://doi.org/10.5281/zenodo.10420396. ANS temperature measurements used in supplemental (Extended Data Fig. 2) were recorded at the Abisko Scientific Research Station (ANS), Sveriges meteorologiska och hydrologiska institut (SMHI). (2024). Meteorological data from Abisko Observatory (Abisko: station 188800), daily mean air temperatures, 2011-2017. Retrieved [05/30/2024], from https://www.smhi.se/data/meteorologi/ladda-ner-meteorologiska-observationer#stationid=188800.

All metagenomes and metatranscriptomes used in this study are available in NCBI under the umbrella BioProject [PRJNA888099](https://www.ncbi.nlm.nih.gov/bioproject/888099). SRA accessions for the reads used for the interannual microbiome comparisons are listed in Supplementary Table 4, and those used to generate the MAGs are listed in the metadata table accompanying those MAGs (https://doi.org/10.5281/zenodo.13901514). Metagenome-assembled genomes (MAGs) are available in Zenodo at https://doi.org/10.5281/zenodo.13901514.

## References and Notes

1. Rantanen, M. et al. The Arctic has warmed nearly four times faster than the globe since 1979. Commun. Earth Environ. 3, 1–10 (2022).

2. Schuur, E. A. G. et al. Permafrost and Climate Change: Carbon Cycle Feedbacks From the Warming Arctic. Annu. Rev. Environ. Resour. 47, 343–371 (2022).

3. Hugelius, G. et al. Estimated stocks of circumpolar permafrost carbon with quantified uncertainty ranges and identified data gaps. Biogeosciences 11, 6573–6593 (2014).

4. Kuhn, M. A. et al. Controls on Stable Methane Isotope Values in Northern Peatlands and Potential Shifts in Values Under Permafrost Thaw Scenarios. J. Geophys. Res. Biogeosciences 129, e2023JG007837 (2024).

5. Varner, R. K. et al. Permafrost thaw driven changes in hydrology and vegetation cover increase trace gas emissions and climate forcing in Stordalen Mire from 1970 to 2014. Philos. Trans. R. Soc. Math. Phys. Eng. Sci. 380, 20210022 (2021).

6. Jansson, J. K. & Hofmockel, K. S. Soil microbiomes and climate change. Nat. Rev. Microbiol. 18, 35–46 (2020).

7. Hough, M. et al. Coupling plant litter quantity to a novel metric for litter quality explains C storage changes in a thawing permafrost peatland. Glob. Change Biol. 28, 950–968 (2022).

8. Persson, A., Hasan, A., Tang, J. & Pilesjö, P. Modelling Flow Routing in Permafrost Landscapes with TWI : An Evaluation against Site-Specific Wetness Measurements. Trans. GIS 16, 701–713 (2012).

9. Woodcroft, B. J. et al. Genome-centric view of carbon processing in thawing permafrost. Nature 560, 49–54 (2018).

10. Wu, L. et al. Reduction of microbial diversity in grassland soil is driven by long-term climate warming. Nat. Microbiol. 7, 1054–1062 (2022).

11. Wilson, R. M., et al. Soil metabolome response to whole-ecosystem warming at the Spruce and Peatland Responses under Changing Environments experiment. Proc. Natl. Acad. Sci. 118, e2004192118 (2021).

12. De Vries, F. & Shade, A. Controls on soil microbial community stability under climate change. Front. Microbiol. 4, (2013).

13. Weedon, J. T. et al. Compositional Stability of the Bacterial Community in a Climate-Sensitive Sub-Arctic Peatland. Front. Microbiol. 8, (2017).

14. Chang, C. & HilleRisLambers, J. Integrating succession and community assembly perspectives. F1000Research 5, 2294 (2016).

15. Fofana, A. et al. Mapping substrate use across a permafrost thaw gradient. Soil Biol. Biochem. 175, 108809 (2022).

16. Shade, A. et al. Fundamentals of Microbial Community Resistance and Resilience. Front. Microbiol. 3, (2012).

17. Prosser, J. I. Ecosystem processes and interactions in a morass of diversity. FEMS Microbiol. Ecol. 81, 507–519 (2012).

18. Curiel Yuste, J., et al. Strong functional stability of soil microbial communities under semiarid Mediterranean conditions and subjected to long-term shifts in baseline precipitation. Soil Biol. Biochem. 69, 223–233 (2014).

19. DeAngelis, K. M. et al. Long-term forest soil warming alters microbial communities in temperate forest soils. Front. Microbiol. 6, (2015).

20. Dove, N. C. et al. Depth dependence of climatic controls on soil microbial community activity and composition. ISME Commun. 1, 1–11 (2021).

21. Feng, J. et al. Warming-induced permafrost thaw exacerbates tundra soil carbon decomposition mediated by microbial community. Microbiome 8, 3 (2020).

22. Gutknecht, J. L. M., Field, C. B. & Balser, T. C. Microbial communities and their responses to simulated global change fluctuate greatly over multiple years. Glob. Change Biol. 18, 2256–2269 (2012).

23. Monteux, S. et al. Long-term in situ permafrost thaw effects on bacterial communities and potential aerobic respiration. ISME J. 12, 2129–2141 (2018).

24. Xue, K. et al. Tundra soil carbon is vulnerable to rapid microbial decomposition under climate warming. *Nat*. Clim. Change 6, 595–600 (2016).

25. Zhu, K., Chiariello, N. R., Tobeck, T., Fukami, T. & Field, C. B. Nonlinear, interacting responses to climate limit grassland production under global change. Proc. Natl. Acad. Sci. 113, 10589–10594 (2016).

26. Kyker-Snowman, E. et al. Increasing the spatial and temporal impact of ecological research: A roadmap for integrating a novel terrestrial process into an Earth system model. Glob. Change Biol. 28, 665–684 (2022).

27. Wieder, W. R., Bonan, G. B. & Allison, S. D. Global soil carbon projections are improved by modelling microbial processes. *Nat*. Clim. Change 3, 909–912 (2013).

28. Emerson, J. B. et al. Host-linked soil viral ecology along a permafrost thaw gradient. Nat. Microbiol. 3, 870–880 (2018).

29. McCalley, C. K. et al. Methane dynamics regulated by microbial community response to permafrost thaw. Nature 514, 478–481 (2014).

30. Mondav, R. et al. Microbial network, phylogenetic diversity and community membership in the active layer across a permafrost thaw gradient. Environ. Microbiol. 19, 3201–3218 (2017).

31. Christensen, T. R. et al. Thawing sub-arctic permafrost: Effects on vegetation and methane emissions. Geophys. Res. Lett. 31, (2004).

32. Hodgkins, S. B. et al. Changes in peat chemistry associated with permafrost thaw increase greenhouse gas production. Proc. Natl. Acad. Sci. 111, 5819–5824 (2014).

33. Beisner, B., Haydon, D. & Cuddington, K. Alternative stable states in ecology. Front. Ecol. Environ. 1, 376–382 (2003).

34. Freire-Zapata, V. et al. Microbiome–metabolite linkages drive greenhouse gas dynamics over a permafrost thaw gradient. Nat. Microbiol. 10.1038/s41564-024-01800-z (2024) doi:10.1038/s41564-024-01800-z.

35. McGivern, B. B. et al. Microbial polyphenol metabolism is part of the thawing permafrost carbon cycle. Nat. Microbiol. 9, 1454–1466 (2024).

36. NOAA National Centers for Environmental Information. NOAA National Centers for Environmental Information, Monthly Global Climate Report for Annual 2017. https://www.ncei.noaa.gov/access/monitoring/monthly-report/global/201713 (2018).

37. Stegen, J. C. et al. Quantifying community assembly processes and identifying features that impose them. ISME J. 7, 2069–2079 (2013).

38. Wilson, R. M. et al. Plant organic matter inputs exert a strong control on soil organic matter decomposition in a thawing permafrost peatland. Sci. Total Environ. 820, 152757 (2022).

39. Du, E. et al. Global patterns of terrestrial nitrogen and phosphorus limitation. Nat. Geosci. 13, 221–226 (2020).

40. Elser, J. J. et al. Global analysis of nitrogen and phosphorus limitation of primary producers in freshwater, marine and terrestrial ecosystems. Ecol. Lett. 10, 1135–1142 (2007).

41. Perryman, C. R. et al. Thaw Transitions and Redox Conditions Drive Methane Oxidation in a Permafrost Peatland. J. Geophys. Res. Biogeosciences 125, e2019JG005526 (2020).

42. Hodgkins, S. B. Changes in organic matter chemistry and methanogenesis due to permafrost thaw in a subarctic peatland. (The Florida State University, United States -- Florida, 2016).

43. Graham, E. B., Yang, F., Bell, S. & Hofmockel, K. S. High Genetic Potential for Proteolytic Decomposition in Northern Peatland Ecosystems. Appl. Environ. Microbiol. 85, (2019).

44. Norman, J. S. & Friesen, M. L. Complex N acquisition by soil diazotrophs: how the ability to release exoenzymes affects N fixation by terrestrial free-living diazotrophs. ISME J. 11, 315–326 (2017).

45. Lerch, B. A., Smith, D. A., Koffel, T., Bagby, S. C. & Abbott, K. C. How public can public goods be? Environmental context shapes the evolutionary ecology of partially private goods. PLOS Comput. Biol. 18, e1010666 (2022).

46. Delmont, T. O. et al. Nitrogen-fixing populations of Planctomycetes and Proteobacteria are abundant in surface ocean metagenomes. Nat. Microbiol. 3, 804–813 (2018).

47. Beier, S. & Bertilsson, S. Bacterial chitin degradation—mechanisms and ecophysiological strategies. Front. Microbiol. 4, (2013).

48. Belova, S. E. et al. Hydrolytic Capabilities as a Key to Environmental Success: Chitinolytic and Cellulolytic Acidobacteria From Acidic Sub-arctic Soils and Boreal Peatlands. Front. Microbiol. 9, (2018).

49. De Tender, C. et al. Peat substrate amended with chitin modulates the N-cycle, siderophore and chitinase responses in the lettuce rhizobiome. Sci. Rep. 9, 9890 (2019).

50. Kaczmarek, M. B., Struszczyk-Swita, K., Li, X., Szczęsna-Antczak, M. & Daroch, M. Enzymatic Modifications of Chitin, Chitosan, and Chitooligosaccharides. Front. Bioeng. Biotechnol. 7, (2019).

51. Lopes, W., Amor, D. R. & Gore, J. Cooperative growth in microbial communities is a driver of multistability. Nat. Commun. 15, 4709 (2024).

52. Canfield, D. E. et al. A Cryptic Sulfur Cycle in Oxygen-Minimum–Zone Waters off the Chilean Coast. Science 330, 1375–1378 (2010).

53. Helbert, W. Marine Polysaccharide Sulfatases. Front. Mar. Sci. 4, (2017).

54. Van Vliet, D. M. et al. Anaerobic Degradation of Sulfated Polysaccharides by Two Novel Kiritimatiellales Strains Isolated From Black Sea Sediment. Front. Microbiol. 10, 253 (2019).

55. Oliverio, A. M. et al. Rendering the metabolic wiring powering wetland soil methane production. Preprint at 10.1101/2024.02.06.579222 (2024).

56. Tveit, A. T., Urich, T., Frenzel, P. & Svenning, M. M. Metabolic and trophic interactions modulate methane production by Arctic peat microbiota in response to warming. Proc. Natl. Acad. Sci. 112, E2507–E2516 (2015).

57. Ciccarese, D. et al. Rare and localized events stabilize microbial community composition and patterns of spatial self-organization in a fluctuating environment. ISME J. 16, 1453–1463 (2022).

58. Embree, M., Liu, J. K., Al-Bassam, M. M. & Zengler, K. Networks of energetic and metabolic interactions define dynamics in microbial communities. Proc. Natl. Acad. Sci. 112, 15450–15455 (2015).

59. Ivanova, A. A., Wegner, C.-E., Kim, Y., Liesack, W. & Dedysh, S. N. Identification of microbial populations driving biopolymer degradation in acidic peatlands by metatranscriptomic analysis. Mol. Ecol. 25, 4818–4835 (2016).

60. Kang, H., Freeman, C., Soon park, S. & Chun, J. N-Acetylglucosaminidase activities in wetlands: a global survey. Hydrobiologia 532, 103–110 (2005).

61. Wörner, S. & Pester, M. Microbial Succession of Anaerobic Chitin Degradation in Freshwater Sediments. Appl. Environ. Microbiol. 85, e00963–19 (2019).

62. Evans, P. N. et al. An evolving view of methane metabolism in the Archaea. Nat. Rev. Microbiol. 17, 219–232 (2019).

63. Sánchez-Andrea, I. et al. The reductive glycine pathway allows autotrophic growth of Desulfovibrio desulfuricans. Nat. Commun. 11, 5090 (2020).

64. Allison, S. D. & Martiny, J. B. H. Resistance, resilience, and redundancy in microbial communities. Proc. Natl. Acad. Sci. 105, 11512–11519 (2008).

65. Doak, D. F. et al. The Statistical Inevitability of Stability-Diversity Relationships in Community Ecology. Am. Nat. 151, 264–276 (1998).

66. Thibaut, L. M. & Connolly, S. R. Understanding diversity–stability relationships: towards a unified model of portfolio effects. Ecol. Lett. 16, 140–150 (2013).

67. Bell, T. H. & Bell, T. Many roads to bacterial generalism. FEMS Microbiol. Ecol. 97, fiaa240 (2020).

68. Guo, J. et al. Mobile genetic elements that shape microbial diversity and functions in thawing permafrost soils. Preprint at 10.1101/2025.02.12.637893 (2025).

69. HilleRisLambers, J., Adler, P. B., Harpole, W. S., Levine, J. M. & Mayfield, M. M. Rethinking Community Assembly through the Lens of Coexistence Theory. Annu. Rev. Ecol. Evol. Syst. 43, 227–248 (2012).

70. Varner, R. K., et al. From Archaea to the atmosphere: remotely sensing Arctic methane. Preprint at 10.1101/2025.02.13.638097 (2025).

71. Chang, K.-Y. et al. Methane Production Pathway Regulated Proximally by Substrate Availability and Distally by Temperature in a High-Latitude Mire Complex. J. Geophys. Res. Biogeosciences 124, 3057–3074 (2019).

72. Wieder, W. R. et al. Explicitly representing soil microbial processes in Earth system models. Glob. Biogeochem. Cycles 29, 1782–1800 (2015).

